# snputils: A High-Performance Python Library for Genetic Variation and Population Structure

**DOI:** 10.64898/2026.02.28.708618

**Authors:** David Bonet, Marçal Comajoan Cara, Míriam Barrabés, Riccardo Smeriglio, Devang Agrawal, Khaled Aounallah, Margarita Geleta, Albert Dominguez Mantes, Christophe Thomassin, Cole Shanks, Edward C. Huang, Marc Franquesa Monés, Aina Luis, Joan Saurina, Maria Perera, Cayetana López, Benet Oriol Sabat, Jordi Abante, Sonia Moreno-Grau, Daniel Mas Montserrat, Alexander G. Ioannidis

**Affiliations:** Department of Biomedical Data Science, Stanford University, Stanford, CA, USA; Genomics Institute, University of California, Santa Cruz, Santa Cruz, CA, USA; Department of Biomedical Sciences, Universitat de Barcelona, Barcelona, Catalonia, Spain; Department of Electrical Engineering and Computer Science, University of California, Berkeley, Berkeley, CA, USA; Munster Technological University, Cork, Ireland; Politecnico di Torino, Turin, Italy; School of Life Sciences, École Polytechnique Fèdèrale de Lausanne, Lausanne, Switzerland; École Polytechnique, Palaiseau, France; Department of Computer Science, Stanford University, Stanford, CA, USA; Department of Signal Theory and Communications, Universitat Politècnica de Catalunya, Barcelona, Catalonia, Spain; Department of Computer Science, University of California, Los Angeles, Los Angeles, CA, USA; Institute of Neurosciences (UBNeuro), Universitat de Barcelona, Barcelona, Catalonia, Spain

**Author notes:** These authors contributed equally (David Bonet, Marçal Comajoan Cara, Míriam Barrabés).

## Abstract

The increasing size and resolution of genomic and population genetic datasets offer unprecedented opportunities to study population structure and uncover the genetic basis of complex traits and diseases. The collection of existing analytical tools, however, is characterized by format incompatibilities, limited functionality, and computational inefficiencies, forcing researchers to construct fragile pipelines that chain together fragmented command-line utilities and ad hoc scripts. These are difficult to maintain, scale, and reproduce. To address such limitations, we present `snputils,` a Python library that unifies high-performance I/O, transformation, and analysis of genotype, ancestry, and phenotypic information within a single framework suitable for biobank-scale research. The library provides efficient tools for essential operations, including querying, cleaning, merging, and statistical analysis. In addition, it offers classical population genetic statistics with optional ancestry-specific masking. An identity-by-descent module supports reading of multiple formats, filtering and ancestry-restricted segment trimming for relatedness and demographic inference. `snputils` also incorporates ancestry-masking and multi-array functionalities for dimensionality reduction methods, as well as efficient implementations of admixture simulation, admixture mapping, and advanced visualization capabilities. With support for the most commonly used file formats, `snputils` integrates smoothly with existing tools and clinical databases. At the same time, its modular and optimized design reduces technical overhead, facilitating reproducible workflows that accelerate discoveries in population genetics, genomic research, and precision medicine. Benchmarking demonstrates a significant reduction in genotype data loading speed compared to existing Python libraries. The open-source library is available at https://github.com/AI-sandbox/snputils, with full documentation and tutorials at snputils.org.

---

The rapid expansion of large-scale genomic datasets, driven by advances in high-throughput sequencing technologies and the emergence of global biobank initiatives, has enabled major breakthroughs across genomics and is reshaping biomedical discovery and clinical translation ^1–9^. Modern human genomic datasets, which profile millions of variants across hundreds of thousands to millions of individuals, have fueled the development of genome-wide association studies ^10–12^, polygenic risk scores ^13–16^, ancestry inference methods ^17–20^, genotype data simulation ^21–24^, demographic reconstruction ^25;26^, and clinical trait mapping ^27–29^. These resources now support the identification of disease mechanisms, prioritization of therapeutic targets, and risk stratification that can inform screening and prevention in diverse populations, while also revealing gaps for underrepresented groups that must be addressed to ensure equitable benefit. As these biobanks become increasingly diverse in their representation of the world’s population variation, scalable packages for analyzing that variation are becoming crucial. However, existing population genetic tools are constrained by persistent computational bottlenecks and heterogeneous file formats that hinder reproducible execution at scale.

Population genetic software solutions remain fragmented, forcing researchers to assemble ad hoc pipelines from multiple tools with narrow scopes, incompatible input and output formats, and limited interoperability. This fragmentation often requires substantial custom code to bridge gaps, increasing the risk of failures and silent data corruption during manual format conversions. As biobank-scale studies grow in size and complexity, there is a critical need for unified, robust, and adaptable computational tools that can alleviate these bottlenecks and sustain the pace of genomic discovery in diverse cohorts. A range of tools addresses individual components of genomic workflows, each with distinct trade-offs in scalability, flexibility, and analytical depth ^30–39^. Command-line utilities such as PLINK1/2^30;31^ and BCFtools ^33^ are widely used for large-scale data processing and statistical analysis, offering high efficiency and support for widely adopted file formats. However, these tools lack native Python interfaces, limiting their integration into interactive, exploratory, or programmable workflows. VCFtools ^32^, while effective for basic variant filtering, exhibits lower scalability to biobank-scale datasets. GCTA^34^ adds complementary functionality, enabling heritability estimation and mixed-model association tests that scale efficiently to biobank cohorts. Frameworks like Hail ^39^ leverage distributed computing via Spark and offer an expressive language for combining multiple steps, such as merging datasets, quality control, and statistical modeling, into a single pipeline. Existing Python-native libraries for genomics are also specialized. For instance, scikit-allel ^35^ integrates with NumPy and Pandas to support versatile array operations for exploratory workflows. For larger-scale analyses, sgkit ^37^ leverages Dask and xarray, providing a scalable framework with modules for global ancestry inference. Other libraries include pgenlib ^31^ for efficient binary I/O, and tools like cyvcf2^38^ and pysnptools ^36^ that provide Pythonic data access with limited built-in analysis.

To address this gap, we developed `snputils`, a Python library that unifies high-performance I/O operations, data transformations, querying, analysis, simulations, and advanced visualization in a unified framework. Supporting all major file formats, the library incorporates novel methods for admixture analysis and offers significant performance gains through algorithmic optimization and optional GPU acceleration. Table 1 summarizes how `snputils` compares with ten established genotype toolkits across key computational and analytical features. Previously, users were faced with a trade-off: command-line tools offer efficiency but lack Python integration and flexibility, while Python-based alternatives vary in scalability and feature coverage. `snputils` addresses these limitations by combining the computational efficiency of traditional command-line tools with the programmatic flexibility of Python-based frameworks, while providing ancestry and phenotype analysis capabilities within a unified environment. The library supports chunked processing and on-demand data loading, enabling analysis of large-scale genomic datasets that may exceed available RAM. Core algorithms are implemented using vectorized operations with NumPy and optimized routines, while I/O optimization strategies including memory-mapped file access patterns enable rapid access to specific genomic regions, sample subsets, or data fields without full dataset materialization. Additionally, the library incorporates optional GPU acceleration for matrix operations and data transformations that deliver substantial speedups for memory-intensive genomic computations. Table 2 summarizes the key features of `snputils`.

**Table 1:**
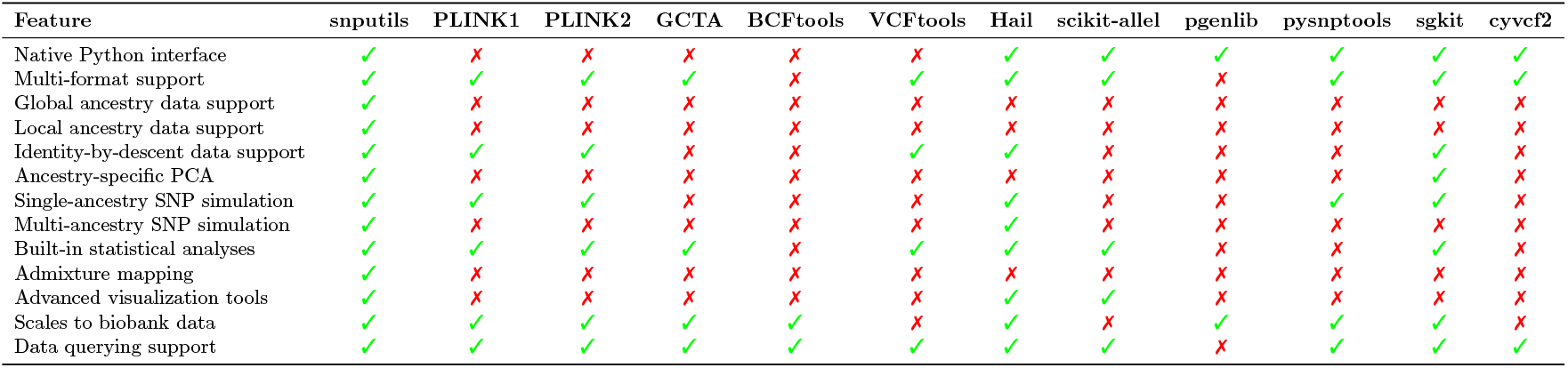
Feature comparison across popular genomic and ancestry analysis tools.

**Table 2:**
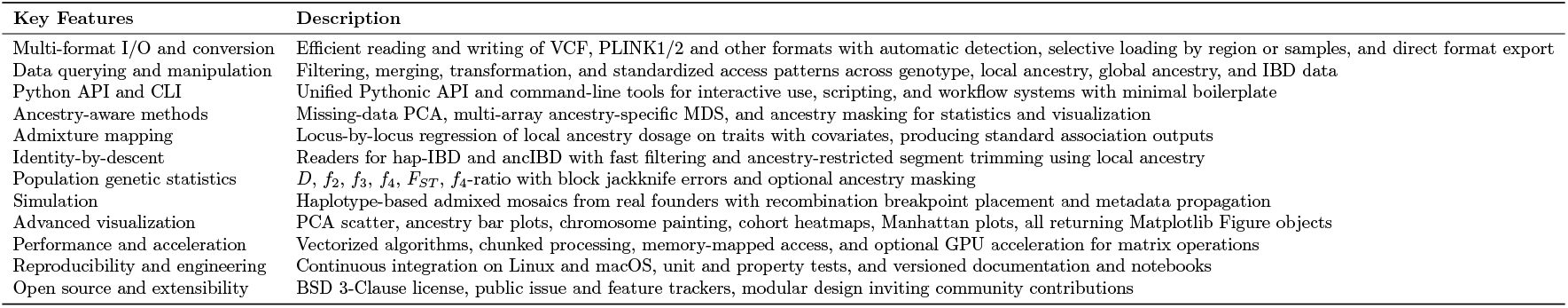
Key features of snputils.

In addition to the Python API, `snputils` provides command-line interface tools that facilitate integration into existing genomic analysis pipelines. These tools enable seamless integration into shell scripts and workflow management systems, supporting both ad hoc data exploration and large-scale automated pipelines.

## Core data model

The `snputils` architecture employs a modular design based on five primary data structures. The `SNPObject` class serves as the primary container for genotype data, implementing efficient storage and manipulation of variant information across samples. Ancestry-related information is managed through two specialized classes: the `LocalAncestryObject,` which stores per-window ancestry assignments for individual samples, and the `GlobalAncestryObject,` which stores per-individual admixture proportions and optional per variant ancestry allele frequencies (P). The unified representation of genotype and ancestry within a shared framework enables researchers to perform integrated analyses that combine population structure assessment with association studies. Additionally, the `PhenotypeObject` provides functionality for both continuous and categorical trait analysis, while the `IBDObject` incorporates identity-by-descent information for relationship and demographic inference. As Python objects, these containers implement native Python interfaces that enable direct interoperability with established scientific computing libraries such as NumPy ^40^, SciPy ^41^, and scikit-learn ^42^, as well as high-performance deep learning libraries including PyTorch ^43^, TensorFlow ^44^, and JAX ^45^. All internal objects support serialization as Python pickle files for efficient storage and reuse.

## File formats and I/O

The library supports the major file formats commonly encountered in population genetics research, providing specialized readers to load data into unified internal structures. Genotype data from VCF files ^32^ (plain or bgzip-compressed), PLINK1^30^ binary filesets (BED/BIM/FAM), PLINK2^31^ formats (PGEN/PVAR/PSAM), and Genotype Representation Graph files (GRG) ^46^ are parsed into `SNPObject` containers; local ancestry files in MSP format compatible with RFMix v2^18^ and G-Nomix ^19^ into `LocalAncestryObject` containers; and admixture proportion matrices (Q/P) produced by ADMIXTURE ^17;47^ and Neural ADMIXTURE ^20^ into `GlobalAncestryObject` containers. Additionally, the library includes corresponding writers for all supported formats, enabling seamless data export and format conversion capabilities. The reader system employs automatic file format detection and provides standardized data access patterns that are independent of underlying file structures. This design eliminates the need for format-specific parsing implementations and reduces potential sources of errors in analytical pipelines. As a minimal example, reading a VCF into a `SNPObject` and extracting the genotype matrix requires only three lines (Listing 1). The readers are optimized for high-throughput data extraction, enabling selective retrieval of specific genomic regions, sample subsets, or data fields without loading entire datasets into memory. Parallelization forms a core component of the optimization approach, with extensive multi-threading implemented across all file readers. The VCF reader specifically uses the Polars library for accelerated parallel processing, while PLINK format readers employ custom parallel algorithms for binary data extraction. Memory-efficient data structures minimize RAM usage during file operations, allowing processing of large-scale datasets that would otherwise exceed system memory limits. The writer system extends these optimizations to output operations, supporting both consolidated multi-chromosome files and parallelized chromosome-specific file generation.

**Listing 1:**
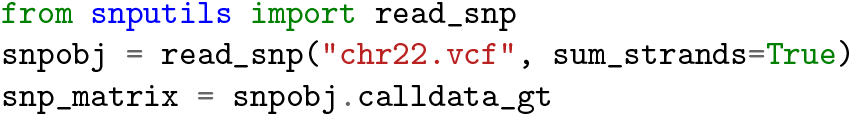
Minimal VCF ingestion with `snputils.`

For identity-by-descent data, `snputils` auto-detects hap-IBD ^48^ and ancIBD ^49^ outputs and parses them into `IBDObject` containers. All parsers are Polars-backed for speed and memory efficiency and standardize fields to a common schema. Manipulation and quality control routines are provided for the different genetic objects, including variant and sample filtering, statistics calculations, among others.

We compared `snputils` readers for BED, PGEN, and VCF with pgenlib ^31^, pysnptools ^36^, sgkit ^37^, scikitallel ^35^, cyvcf2^38^, plinkio ^50^, pandas-plink ^51^, and pysam ^52^ using chromosome 22 data from the 1000 Genomes Project^53^. For VCF, the reported timing corresponds to the minimal read operation in Listing 1 (i.e., the wall-clock time to ingest the file and materialize the genotype matrix), with analogous minimal ingestion for each comparison library^1^. As shown in Figure 1, `snputils` consistently outperformed all alternative tools across all supported formats. For PLINK binary formats, `snputils` achieves reading times of 0.95 seconds for BED files and 0.91 seconds for PGEN files, representing performance improvements of up to 99.91% compared to existing Python-based tools that require between 13 seconds and up to several minutes for equivalent operations. VCF file processing demonstrates even more substantial gains, with `snputils` completing reads in 1.48 seconds compared to 39-68 minutes required by alternative implementations, representing up to 97.38% reduction in processing time.

**Figure 1.**
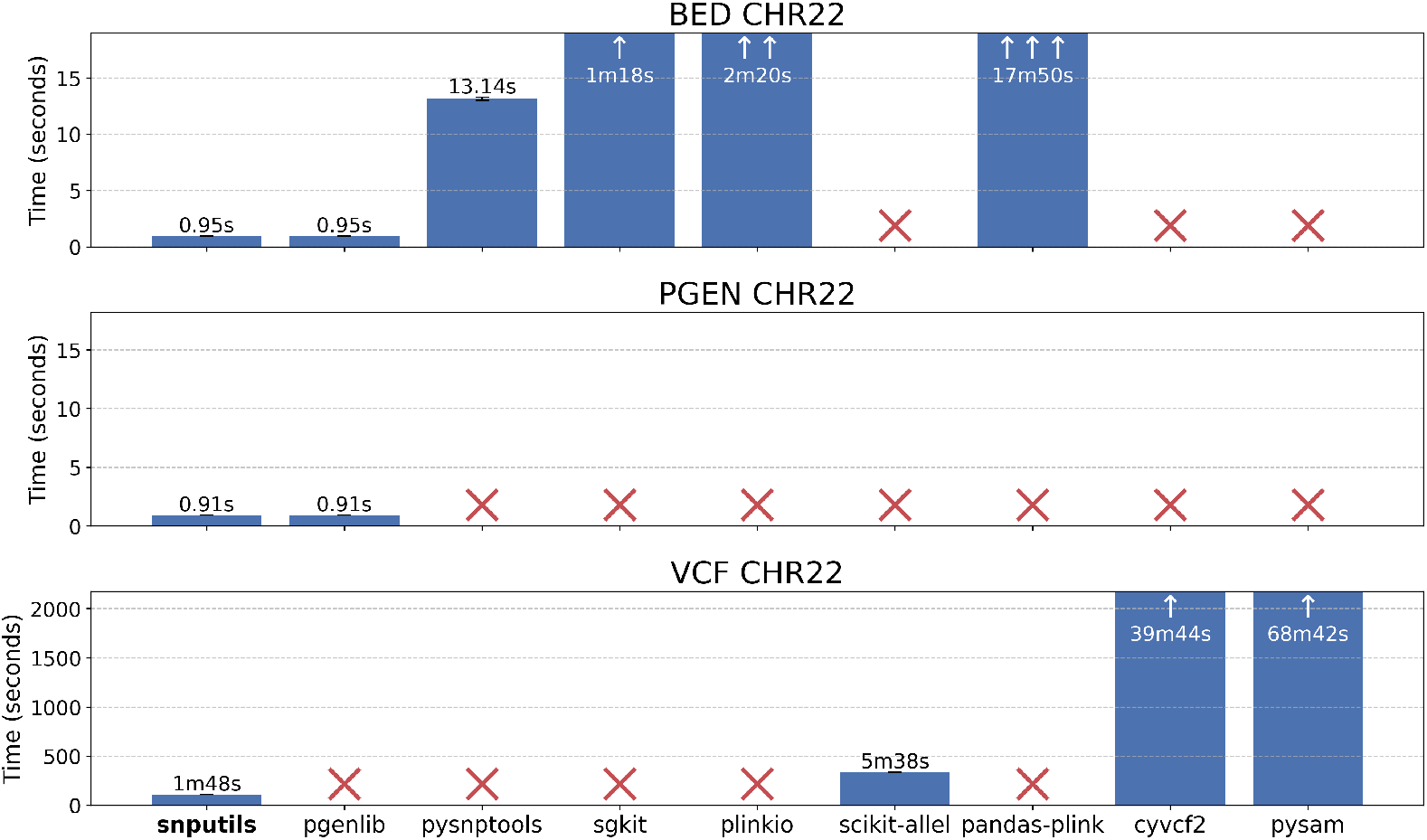
Benchmark reading time comparison across formats and tools for reading chromosome 22 SNPs of all the samples in the 1000 Genomes Project ^53^.

Beyond CPU-based optimizations, `snputils` incorporates optional GPU acceleration through PyTorch integration to improve computational performance. The TorchPCA implementation leverages CUDA-enabled computation for principal component analysis, providing substantial performance improvements over CPU-based alternatives that are particularly beneficial for large-scale population genetics studies where computational efficiency directly impacts analytical feasibility. This GPU acceleration delivers 3-5x speedups for downstream matrix algebra operations compared to CPU-only implementations.

## Simulation

`snputils` includes a lightweight haplotype-based simulator that generates mosaics of haplotypes from real, phased founder haplotypes. Given a VCF, per-sample metadata, and optionally a genetic map, the simulator samples a generation-since-admixture parameter per batch. It then places recombination breakpoints along each chromosome, with probabilities proportional to the local genetic distance if a map is supplied (or uniformly otherwise). At each breakpoint, it switches the donor haplotype by permuting suffix segments across the founder pool, which yields tract length distributions that scale with genetic length and the generation parameter. Both discrete labels (e.g., population codes) and continuous labels (e.g., latitude and longitude, optionally stored as 3D unit vectors) are propagated through the mosaic and can be aggregated per SNP window by the mean or by an approximate GPU-friendly mode. This “select and stitch” approach is consistent with the standard Poisson model of crossovers used in prior simulation work ^54^, while remaining simple to run at scale through a CLI that batches simulations and writes NumPy archives or `SNPObject` pickles.

## Statistics

`snputils` provides a unified suite of population genetic statistics, including *D* ^55^, *f*_2_, *f*_3_, *f*_4_, the *f*_4_-ratio ^56^, and the fixation index (*F*_*ST*_) with support for both Hudson ^57^ and Weir-Cockerham ^58^ methods. Each estimator is computed from per-SNP contributions aggregated over blocks, and accompanied by delete-one block jackknife standard errors ^59^. Inputs can be either aggregated allele frequency and count matrices with population names or a `SNPObject` plus per-sample labels, which are internally converted to population-level allele frequencies. All statistics support ancestry-specific analyses (see below) by masking genotypes to a chosen local ancestry using a `LocalAncestryObject` before aggregation, which enables ancestry-stratified tests without pre-splitting datasets. The routines are vectorized, tolerant to missingness, and scale to biobank data.

## Ancestry and admixture analyses

The library supports various population genetic methods and data types related to genetic ancestry analysis. These include parsers for global and local ancestry inference outputs, alongside downstream tools like ancestry masking for PCA, ancestry-specific computation of allele statistics, and efficient admixture mapping.

Global ancestry proportions from Q and P format files are supported.The library supports merging multiple runs and visualizing results as ordered bar plots with samples grouped by metadata and colored palettes for easy interpretation of population composition. Local ancestry painting functions render colored tracts along chromosomes for individual samples and generate cohort-level heatmaps of ancestry frequencies in genomic windows. These visualizations facilitate detailed examination of fine-scale admixture patterns and tract distributions across populations. These capabilities integrate with the F-statistics module, which can optionally restrict computations to a specified local ancestry to test hypotheses within ancestry components.

A complete admixture mapping pipeline automates locus-by-locus regression of local ancestry dosage against quantitative or binary traits, controlling for global ancestry PCs and other covariates. The pipeline accepts local ancestry calls and phenotype tables and produces standard association files. Results output as PLINK-format association files.

## Identity-by-descent and relatedness

`snputils` includes a unified identity-by-descent module that reads hap-IBD ^48^ and ancIBD ^49^ (for analysis of ancient samples) outputs into a common `IBDObject` representation, with automatic format detection and gzip support. Convenience methods support fast filtering by chromosome, individuals, segment type, and minimum length. For ancestry-aware analyses in admixed cohorts, ancestry masking can be performed by intersecting with a `LocalAncestryObject` to isolate IBD segments within intervals where both individuals carry a specified local ancestry. The restriction is haplotype-aware when haplotype IDs are present and can optionally require both haplotypes to match the target ancestry. Segments are either entirely filtered out or subsegments are clipped to ancestry window boundaries and returned with cM lengths computed from genetic positions when available. These capabilities enable ancestry-specific relatedness screening and demographic inference and integrate naturally with downstream admixture and association workflows.

## Design and Implementation

The `snputils` library adheres to modern software engineering practices, with comprehensive dependency management and testing frameworks. The library supports maintained Python releases, with continuous integration that tests Linux and macOS builds and unit and property tests covering readers, transformations, statistics, and plotting. Continuous integration workflows ensure code quality and cross-platform compatibility, while comprehensive documentation and example notebooks that facilitate user adoption are versioned with releases, and public issue and feature-request trackers support community feedback. The open-source nature of the project, distributed under a BSD 3-Clause License, promotes community adoption and contribution. Comprehensive documentation, example notebooks, and active issue tracking facilitate user support and community engagement.

## Visualization

Users can generate scatter plots for principal component analyses, admixture proportions bar plots, chromosome paintings at both sample and cohort levels, Manhattan plots for genome-wide association results, and more. All plotting functions are implemented as methods on the core Python objects described and return Matplotlib Figure objects, allowing full customization of color palettes, annotation layers, and layout via a consistent interface. The toolkit avoids external plotting dependencies to ensure consistent figure rendering across environments and supports both interactive notebook workflows and scripted analyses.

In summary, `snputils` unifies genotype, ancestry, phenotype, and identity-by-descent analyses in a single Python library that scales to biobank data. It combines efficient I/O and GPU-ready computation with ancestry-aware methods such as multi-array ancestry-specific MDS, and an admixture mapping pipeline. The API reduces intermediary code and improves reproducibility for interactive and pipeline use in population genetics and biobank analyses. The `snputils` library is a contribution to the genomic analysis software ecosystem, addressing critical needs for performance, usability, and methodological innovation. Its comprehensive approach to genomic data handling, combined with novel analytical methods and high-performance implementations, positions it as a user-friendly and effective tool for contemporary population genetics and genomic studies. The library’s emphasis on both computational efficiency and methodological rigor makes it particularly suitable for large-scale genomic analyses where traditional approaches may prove computationally prohibitive or methodologically insufficient. The benchmarks demonstrate significant computational speed improvements, which are particularly relevant for large-scale genomic studies.

By consolidating multiple functionalities into a single Python library, `snputils` fosters a streamlined workflow for population genetics applications. Its efficient and intuitive interfaces reduce barriers to large-scale genetic analyses, encouraging reproducibility and collaboration. Future developments include additional support for emerging file formats. The software and documentation are freely available at snputils.org. We encourage the community to adopt, test, and contribute to `snputils` for future genetic studies, clinical analyses, and large-scale cohort investigations.

## Web resources

Documentation and tutorials for `snputils` are available at snputils.org, and the software is available as `snputils` on PyPI.

## Data and Code Availability

Code: https://github.com/AI-sandbox/snputils.

Data for the benchmarks (1KGP^53^): https://ftp.1000genomes.ebi.ac.uk/vol1/ftp/data_collections/1000_genomes_project/release/20181203_biallelic_SNV.

## Footnotes

1 Benchmark code available in https://github.com/AI-sandbox/snputils/tree/main/benchmark

